# Urbanisation impacts plumage colouration in a songbird across Europe: evidence from a correlational, experimental, and meta-analytical approach

**DOI:** 10.1101/2022.09.13.507844

**Authors:** Pablo Salmón, David López-Idiáquez, Pablo Capilla-Lasheras, Javier Pérez-Tris, Caroline Isaksson, Hannah Watson

## Abstract

Urbanisation is increasing at a phenomenal rate across the globe, transforming landscapes, presenting organisms with novel challenges, shaping phenotypic traits, and even impacting fitness. Among colour traits, urban individuals are widely claimed to have duller tones in carotenoid-based traits, the so-called “*urban dullness*” phenomenon. However, at the intra-specific level, this generalisation is surprisingly inconsistent and often based on examples from single urban/non-urban population pairs or a limited geographic area. Here, combining correlational, experimental, and meta-analytical results from a common songbird, the great tit (*Parus major*), we investigated carotenoid-based plumage coloration in urban and forest populations across Europe. We find that, as predicted, urban individuals are paler than forest individuals. Interestingly, we also find large population-specific differences in the magnitude of the urban-forest contrast in plumage colouration. Moreover, our meta-analysis indicates a non-significant effect of environmental pollution on carotenoid-based plumage for the species, suggesting that the observed differences across urban populations are not only driven by pollution. Finally, using one region as an example (Malmö, Sweden), we reveal population-specific processes behind plumage colouration differences, which are likely the result of variation in the spatial and temporal distribution of carotenoid-rich resources in anthropogenic environments. This is the first study to quantify the consistency of an oft-cited textbook example of the impact of urbanisation on wildlife; our results provide the most convincing evidence to date of the “*urban dullness*” phenomenon, but also highlight that the magnitude of the phenomenon depends on local urban characteristics.

## 1 Introduction

The human urban population is growing exponentially in parallel with the global expansion of urban development (Seto et al., 2012; United Nations, 2018). This unprecedented increase in urban landscapes strongly affects the environmental conditions encountered by organisms and poses a major challenge for wildlife across the globe (Hendry et al., 2017). Despite urban habitats being relatively young ecosystems, their novel selective pressures can dramatically shape the phenotypic and genotypic variation of populations living in them (e.g., Salmón et al., 2021; Santangelo et al., 2022; Thompson et al., 2022). In recent decades, research across the globe has demonstrated divergence in multiple phenotypic traits between urban and non-urban populations, across a variety of taxa (Alberti et al., 2017; Sepp et al., 2018; Capilla-Lasheras et al., 2022). Even though one of the most well-known examples of natural selection on a phenotypic trait arose due to urban industrial pollution - the case of the peppered moth (Steward, 1977) - animal coloration has been surprisingly little studied in the context of urbanisation (reviewed in Leveau, 2021). Colour traits play significant roles in sexual signalling and camouflage and can therefore strongly influence survival and reproductive success. The evidence, to date, suggests that urban organisms tend to present higher levels of melanisation (“*urban melanism*”) and duller carotenoid-based traits (“*urban dullness*”). However, these results are largely based on examples from one or a few populations and/or a limited geographical area (Leveau, 2021), and there has been no attempt yet to comprehensively evaluate the extent to which these effects can be generalised across larger spatial scales. Given that the scale and attributes of urban environmental change can strongly depend on local characteristics (Alberti et al., 2020), it is especially important to understand the uniformity of widely asserted urban impacts.

Carotenoid-based colouration (i.e., yellow, orange, and red colours) is central to widespread and conspicuous ornamental traits in animals (Blount & McGraw, 2008) and is often shown to be under strong sexual selection (Svensson & Wong, 2011). Carotenoid-based colouration has received much attention in the literature, particularly because it is strongly affected by environmental conditions (e.g., Fitze et al., 2009; Martínez-Padilla et al., 2007; Ruell et al., 2013) and is potentially linked to individual quality (e.g., Weaver et al., 2018). Animals cannot synthesise carotenoids *de novo* and must obtain them from the diet (Goodwin, 1986).

Aspects of the urban environment, such as air pollution and the predominance of non-native tree planting in urban parks, could alter animal carotenoid colouration via changes in the bioavailability of carotenoids in the diet (Eeva et al., 1998; Sillanpää et al., 2008). For example, primary producers and the caterpillars that feed upon them have been shown to have lower carotenoid levels in polluted, compared with unpolluted, areas (Silanpää et al. 2008), and urban-dwelling caterpillars contain less carotenoids than those in the forest (Isaksson & Andersson, 2007). Caterpillars are the main source of carotenoids deposited in the plumage of several songbird species, including the great tit (*Parus major*). Indeed, the great tit - a model system in avian urban ecology and common across Europe - is known to exhibit *duller* plumage colouration in some urban populations (Biard et al., 2017; Hõrak et al., 2000; Isaksson et al., 2005), probably reflecting carotenoid-deficient diets. Additionally, carotenoids are often regarded as key immunomodulators and antioxidants (Pérez-Rodríguez, 2009; but see e.g., Koch et al., 2018); thus, observed differences in carotenoid-based colouration between urban and non-urban populations might also be the result of trade-offs between ornamentation and health, driven by increased pathogen exposure or higher levels of oxidative stress in urban environments (e.g., Bortolotti et al., 2003; Isaksson et al., 2005; Isaksson & Andersson, 2008).

Despite increasing interest in urban ecology research, in recent decades, our current knowledge of the impact of urbanisation on animal colour traits is based on a few single populations, even for ecologically well-studied species (Leveau, 2021). Moreover, studies often use a single methodological approach, and no study to date have combined correlative data at a large geographical scale, detailed field experiments and research synthesis to provide a holistic picture of how urbanisation affects organismal colouration. These limitations hinder our ability to generalise about urban effects on colouration across larger spatial and temporal scales. For instance, methodological differences across studied populations might increase the between-site variance, but also the key factors influencing carotenoid-based traits, such as food availability or plant diversity and abundance, are expected to differ locally across urban environments (Szulkin et al., 2020).

Here, we aim to quantify, for the first time, the spatial uniformity of the “*urban dullness*” phenomenon to gain understanding of the possible underlying causes. Using a case study of the great tit, an ecologically relevant avian model in Europe whose characteristic yellow feathers are a well-known example of a carotenoid-based ornament (Broggi & Senar, 2009; Evans & Sheldon, 2012, 2013; Gamero et al., 2015), we combine complementary approaches, utilising correlational data at a continental scale, a cross-fostering experiment and a meta-analysis. Previous work in this widely distributed bird species within Eurasia has shown dramatic phenotypic shifts in response to urbanisation across multiple traits (e.g., Branston et al., 2021; Caizergues et al., 2018; Charmantier et al., 2017; Corsini et al., 2017; Hardman & Dalesman, 2018; Isaksson et al., 2009; Senar et al., 2017; Sprau et al., 2017), including carotenoid-based colouration and its physiological basis (see e.g., Biard et al., 2017; Hõrak et al., 2000, in Table S2). However, other studies have also found no differences in carotenoid-based plumage traits (e.g., Isaksson et al. 2007), suggesting an inconsistency in the pattern.

Firstly, we test if urbanisation is consistently correlated with breast carotenoid-based colouration in adult birds, using a paired sampling design of urban and forest populations across five urban centres in Europe (Figure 1a). Secondly, in a reciprocal cross-fostering experiment within one of the population pairs (Malmö, Sweden), we quantify the relative contribution of environmental *versus* genetic/parental influence on colour variation of chicks. Carotenoid-based plumage in great tits is fully established in the second year of life, and it is highly repeatable across moult cycles thereafter (Evans & Sheldon, 2013). However, little is known about how urban habitats might influence the acquisition of adult plumage colouration. Therefore, thirdly, again using focal urban and forest populations in Malmö, we also investigate how colouration varies with age in relation to urbanisation. Finally, ecotoxicology studies suggest a link between certain urban stressors, such as pollutants, and great tit colouration (Eeva et al., 1998). To better understand the overall effect of urbanisation on colouration, fourthly and finally, we conducted a meta-analysis of the literature concerning changes in breast colour in response to urbanisation and pollution in great tits. Overall, our multi-analytical approach provides a comprehensive understanding of the repeatability of the “*urban dullness*” phenomenon across a large spatial scale and investigate possible driving mechanisms.

**Figure 1.**
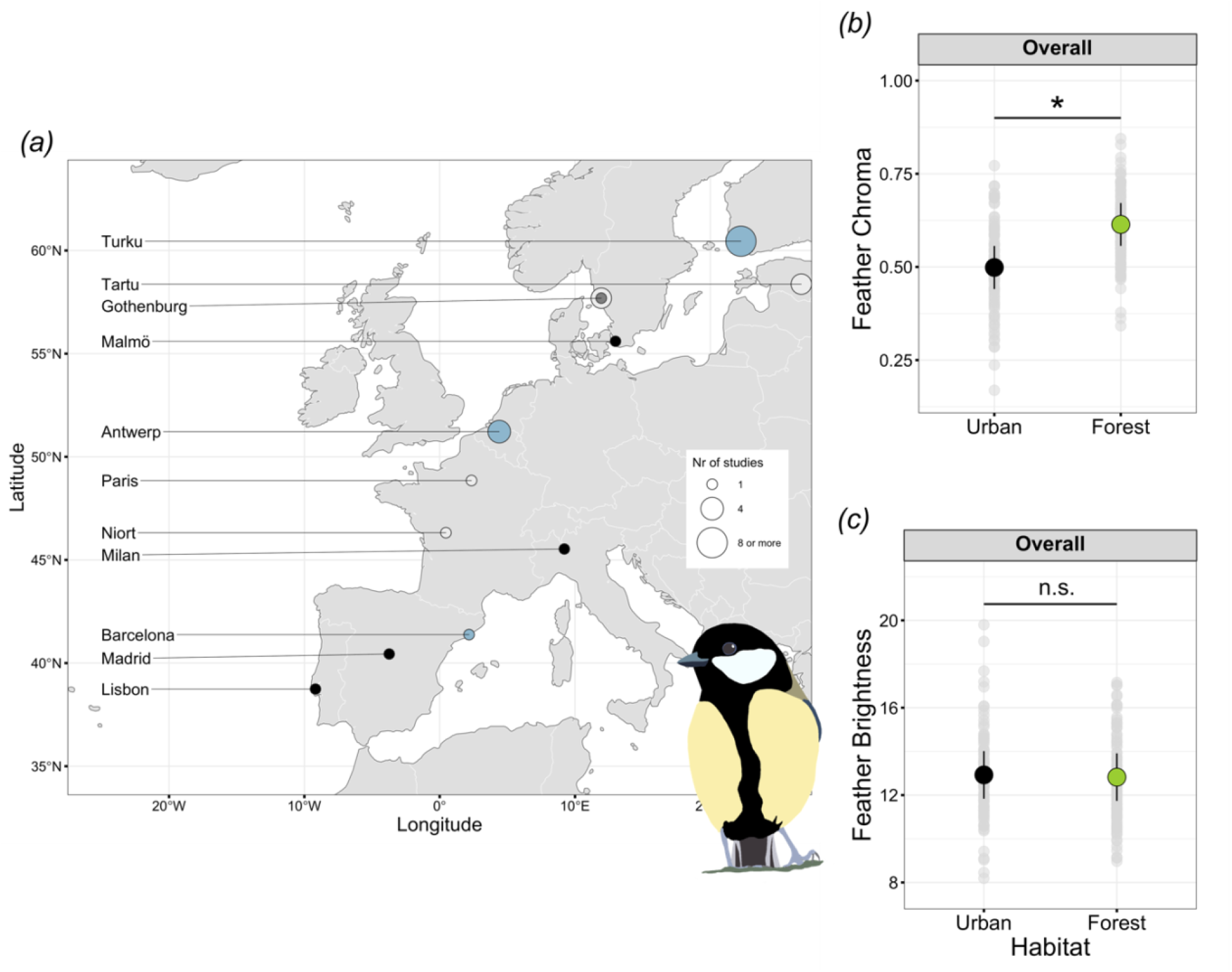
(*a*) Locations of great tit (*Parus major*) populations across Europe included in this study. Black circles: urban/forest population pairs sampled in the empirical study in which differences in breast plumage colour traits were compared (see Table S1 for details); white and blue circles: populations from the literature on effects of urbanisation (white) or pollution (blue) on colour traits and included in the meta-analysis (see Table S2 for details). For adult birds (1 yr and 2+ yr) sampled in the empirical study, i.e., black circles in (a): *(b)* chroma reflects the amount of pigment in the feathers (i.e., carotenoids) and *(c)* brightness reflects the structural quality of feathers. Sample sizes (urban/forest): Gothenburg (11/11), Malmö (42/41), Milan (16/18), Madrid (16/20), Lisbon (18/16). Asterisks denote significant differences between urban and forest habitats with p <0.05 (habitat main term see Table S4). n.s. = non-significant differences. Model mean values (solid-coloured circles) with 95% confidence intervals are plotted, along with raw data (grey circles).

## 2 Material and Methods

### 2.1 General fieldwork and cross-fostering design

During the years 2014 and 2015 (breeding, i.e., spring, and post breeding, i.e., summer, seasons), we captured a total of 209 adult great tits (aged 1 yr and 2+ yr) in five paired urban-forest habitats across Europe (n= 5 urban /5 forest populations; Figure 1a; see Salmón et al 2021; Table S1). Adult individuals were captured using mist nets (Lisbon, Madrid, and Milan) or caught in nest-boxes during breeding (Gothenburg and Malmö; see Table S1). In all cases, we sampled each urban and forest pair within the same season and year. All urban sites were located in built-up areas or parks within city boundaries (see Salmón et al 2021 for details about Gothenburg, Milan, Madrid and Lisbon, and Andersson et al 2015, Salmón et al 2016, Salmón et al 2017 for details about Malmö). Forest locations were natural/semi-natural forests in areas of very low human habitation. Each pair of study locations were separated by 33.6 ± 5.37 km, a distance far exceeding the mean adult and natal dispersal distance of this species (mean ± SD: adult, 2.5 ± 12.3 km and natal, 5.3 ± 17.9 km (Paradis et al., 1998)).

In 2013, we performed a reciprocal cross-fostering of nestlings between the urban and forest populations in Malmö (see details in Salmón et al., 2016). Briefly, when nestlings were 2 d old (hatching day = 0 d), we exchanged half of the nestlings from a brood in the urban habitat with the same number of nestlings from a nest of the same age and similar brood size (± 1 nestling) in the forest study population (*n* = 10 nest pairs: 73 nestlings). Where brood sizes differed, we swapped the number of chicks corresponding to half the number of the smaller brood. Cross-fostered chicks were marked by clipping the outermost tip of their claws. Nestlings were individually ringed, and breast feathers were collected at 15 d.

Feathers were collected from birds using a standardised methodology, in which a total of eight yellow feathers were pulled from both sides of the upper part of the breast and stored in a paper envelope in a dark place until measurement. These represent the feathers grown during the nestling period (nestlings), autumn post-juvenile partial moult (1 yr) and autumn post-breeding complete moult (2+ yr). Birds were sexed either visually (adults) or molecularly, using primers P2 and P8 (nestlings; Griffiths, Double, Orr, & Dawson, 1998). There was no difference in sex ratio in the dataset between habitats across population pairs (urban-forest): adults (GLMM, χ^2^_1_ = 0.04, p= 0.842), nestlings (GLMM, χ^2^_1_ = 0.13, p= 0.712). For adults, only feathers from individuals whose age category could be determined based on plumage characteristics were used in this study, following L. Svensson (1992). The collection of feather samples was approved by the relevant local regulatory bodies: Malmö-Lund animal Ethical Committee (permit no. M454 12:1), the CEMPA and Portuguese Ministry of Environment (permit nos. 40/2014 and 164/2014), the Ministry of the Environment, Housing and Territorial Planning of Madrid (permit nos. 10/103329.9/14, 10/169940.9/13, 10/045383.9/14, 10/127641.9/14 and 10/055393.9/14) and the Institute for Environmental Protection and Research (ISPRA, license nos.15510 and 15944) and the Lombardy Region (permit no. 3462).

### 2.2 Feather colouration measurements

Feather colouration was measured by a single person (D.L-I) using a spectrophotometer (AVASPEC-2048, Avantes BV, Apeldoorn, Netherlands) and a deuterium-halogen light source (AVALIGHT-DH-S lamp, Avantes BV) covering the range 300-700 nm and kept at a constant angle of 90° from the feathers (Fargevieille et al., 2017). For each bird, we computed the average of six measures of reflectance spectra, with three spectra measured for each of two samples consisting of four breast feathers. Following previous studies (e.g., Lopez-Idiaquez et al., 2022), we computed chromatic and achromatic colour variables based on the shape of the spectra using the R package *pavo* (Maia et al., 2019). Specifically, we computed yellow chroma as (R700-R450)/R700 (hereafter chroma), with higher values of chroma being linked to higher carotenoid content in plumage (Isaksson, McLaughlin, et al., 2007). In addition, we also calculated brightness (area under the reflectance curve divided by the width of the interval at 300-700nm). To estimate the technical repeatability of our colouration measurements, we repeated measurements on a subset of 30 adults (~50:50 1 yr and 2+ yr) and 15 nestlings twice (on different days). A repeatability analysis, using the R package *rptR* (Stoffel et al., 2017), indicates that our measures were highly repeatable for both colour traits and age groups (yellow chroma, adults: R= 0.85, 95%CI [0.697, 0.930]; nestlings: R= 0.79, 95%CI [0.525, 0.924]; yellow brightness, adults: R= 0.87, 95%CI [0.755, 0.939]; nestlings: R= 0.90, 95%CI [0.731, 0.965]; all p<0.001).

### 2.3 Statistical analysis

Firstly, we first investigated differences in brightness and chroma in adult birds from the five urban regions using linear mixed models (LMMs). Models included habitat (urban or forest), sex, age (1 yr or 2+ yr) and all 2-way interactions as fixed effects. Sampling location (n= 10) and urban region (n= 5) were included as random effects. The season when the feathers were collected, i.e., breeding, or post-breeding (see Table S1), did not influence chroma (F_1, 7.99_= 0.011, p= 0.920) nor brightness (F_1, 7.69_= 1.30, p= 0.288); thus, season was not included in the models to avoid overparameterization. In addition, we conducted two separate *ad hoc* LMMs for urban or forest populations, respectively, to explore the habitat variation in chroma colouration across- (percentage explained by the region ID) and within-populations (percentage explained by the residual variance). These models had the same structure as the initial ones but only with urban region as a random effect. Secondly, using LMMs, we analysed the environmental (i.e., habitat - urban or forest-) and early maternal/genetic effects on chroma and brightness in nestlings from the cross-fostering experiment. Models included the rearing habitat (urban or forest), habitat of origin (urban or forest) and their interaction, together with sex as fixed effects. As our experimental broods included nestlings of mixed origin (i.e., partial cross-fostering), we included the nest of origin (to account for genetic and maternal effects) and the nest of rearing (to account for the effects of the common environment) as random effects. Thirdly, to further investigate the age-dependent variation in colouration between urban and forest habitats, we used LMMs to cross-sectionally analyse the habitat effect using one focal urban region (Malmö). We analysed variation in chroma and brightness across three age categories: nestlings, 1 yr and 2+ yr. Data were included from two consecutive years (2013 and 2014) and collected during breeding. We only included birds ringed as nestlings in the urban habitat (i.e., only those of confirmed urban origin), to understand how the environmental conditions during the first (nestling to 1 yr) and subsequent (1 yr to 2+ yr) moult cycles might have influenced individual colouration. Models included habitat, sex, age and all 2-way interactions as fixed effects and territory (i.e., nest-box) as random effect.

All statistical analyses were performed in R 3.5.2 (R Core Team, 2021). The significance of model terms was estimated using F-tests based on Satterthwaite approximation for the denominator degrees of freedom. In all cases, the distribution of model residuals was inspected visually and did not show marked deviations from normality. Significant 2-way interactions were further dissected using pairwise planned comparisons adjusted by Tukey HSD in *emmeans* (Lenth et al., 2018). The explanatory power of the models was calculated as marginal (R^2^_m_) and conditional (R^2^_c_) values using the R package *r2glmm* (Jaeger et al., 2017). LMMs were fitted using the *lme4* R package (Bates et al., 2014).

### 2.3 Meta-analysis: literature search, data extraction and effect size calculation

To synthesise and assess the current knowledge of urbanisation and pollution on coloration in great tits, we systematically searched the peer-reviewed scientific literature. We searched six databases: A&HCI 1975-present; BKCI-S 2005-present; BKCI-SSH 1990-present; ESCI 2015-present; SCI-EXPANDED 1900-present; SSCI 1900-present, within Web of Science (search conducted on August 2^nd^, 2021) using the following search string: TS = (‘urban*’ OR ‘pollut*’) AND (‘color*’ OR ‘colour*’ OR ‘carot*’) AND (‘Parus major’ OR ‘great tit’ OR ‘tit’). This search produced 81 papers, which were complemented by five studies identified as potentially suitable by the authors and estimates from the new empirical data presented in this paper (Figure S1). We read the title and abstract of these 86 papers to determine their suitability for further inspection and inclusion in this meta-analysis. Papers were included in our meta-analysis if they reported the effect of an anthropogenic disturbance (i.e., urbanisation or chemical pollution) on traits related to yellow colouration of great tit plumage (e.g., brightness, chroma or hue; see full details in Tables S2 and S3). We also included studies on urbanisation and pollution that measured carotenoid levels in feathers or plasma as these have been shown to reflect the plumage colouration of interest (Isaksson et al., 2008). We did not include studies that assessed habitat differences between natural/semi-natural populations (i.e., non-urban linked). This resulted in the inclusion of a total of 22 studies (23 studies including the data presented in the current study) and 128 effect sizes in the meta-analysis. All effect sizes were extracted by one author, PS. We extracted correlation coefficients between pollutants and carotenoid-based colour (n = 8 studies; k = 44 effect sizes) and mean values for population comparisons (either urban/forest or polluted/control; n = 20 studies; k = 84 effect sizes). In the latter case, we calculated point-biserial correlation coefficients (Tate, 1954) as a standardised effect size for comparisons between urban and non-urban populations (Kim et al., 2021). Standardised effect sizes (*r*) and their sampling variances were calculated using the R function “*escalc*” in the R package *metafor* (v. 3.0-2; (Viechtbauer, 2010)).

### 2.4 Meta-analysis: statistical analysis and detection of publication bias

We performed multilevel meta-analytic mixed-effect models using the package *metafor* v3.0-2 (Viechtbauer, 2010) in R v.4.1.1. (R Core Team, 2021). In each model, we used the standardised effect size (*r*) as the response variable. We first fitted an intercept-only model to estimate the overall mean effect (i.e., meta-analytic mean) for the association between anthropogenic disturbance (i.e., urbanisation and pollution) and all the great tit yellow breast colouration traits obtained from the literature search (see *Section 2.3*). We then built individual meta-analytic models to investigate potential drivers of differences in coloration between urban and non-urban populations using only the effect sizes of the same colouration traits as evaluated in the present study, i.e., chroma and brightness, together with carotenoid concentration. Models aimed to evaluate the effect of our chosen moderators (see Table S3 for details) on the association between carotenoid-based colouration and anthropogenic disturbance, based on: *(i)* the type of plumage trait, i.e., chroma, brightness or carotenoid concentration (feathers and plasma); *(ii)* the developmental stage, i.e., adult or nestling; and *(iii)* the type of anthropogenic disturbance, i.e., urbanisation or pollution. In all models, we included study geographical location and study ID as random effects, as well as a term for residual variance. We present model estimates with their 95% confidence intervals (CI) throughout. We calculated total heterogeneity (i.e., the total amount of variation after accounting for sampling variance, *I^2^*_total_), the amount of among-location variation (*I^2^*_location_ ; Figure 1a), the amount of among-study variation (*I^2^*_study_) and amount of residual variation (*I^2^*_residual_) using the function the R function “*i2_ml”* in the *metafor* R package (Viechtbauer, 2010). We also estimated the percentage of variation explained by each moderator in our models using R^2^_marginal_ (Nakagawa & Schielzeth, 2013).

To check for possible publication bias, we performed an extra multilevel meta-regression with the same random structure as above but using as moderators *i)* sampling variance (for small-study effects; see Equation 21 in Nakagawa et al., 2022); and *ii)* mean-centred year of study publication to test time-lag bias (Koricheva & Kulinskaya, 2019).

## 3 Results

### 3.1 Adults: urbanisation and colouration across Europe

Urban and forest adult great tits differed in the chroma of the yellow breast colouration, but not in brightness, across the five studied urban regions (Figure 1b and c; Figure S2; Table S4). Adult urban birds had consistently lower chroma than forest birds (Figure 1b). However, the magnitude of the negative urban effect on chroma varied across cities (Figure S2a). The *ad hoc* analysis treating urban and forest populations in separate models reinforced this finding. Among-population variance in chroma was low in urban populations (0.08 units; only 8% of the variation in chroma was explained by differences among urban populations) and much lower than the among-population variance in forest populations (0.43 units; 40% of the variation in chroma was explained by differences among forest populations). In contrast, within-population variance in chroma was higher in urban populations (0.89 units; 92% of the total variation in chroma was within urban population variation) than in forest populations (0.65 units; 60% of the total variation in chroma was within forest population variation)

### 3.2 Nestlings: urbanisation and colouration in a cross-fostering experiment

In contrast to our findings for adults, great tit nestlings reared in the urban habitat did not differ in yellow breast colouration (chroma or brightness) from their forest counterparts in the Malmö population pair (Figure 2a and b; “*Habitat of rearing”* in Table S5). There were no differences in colour traits between cross-fostered and non-cross fostered nestlings, indicating neither habitat of origin nor habitat of rearing influenced colouration (Table S5). Our results indicate that a high proportion of colour variation among nestlings was explained by the family of rearing independently from the rearing habitat *per se*, in particular for chroma (71% of the total variation).

**Figure 2.**
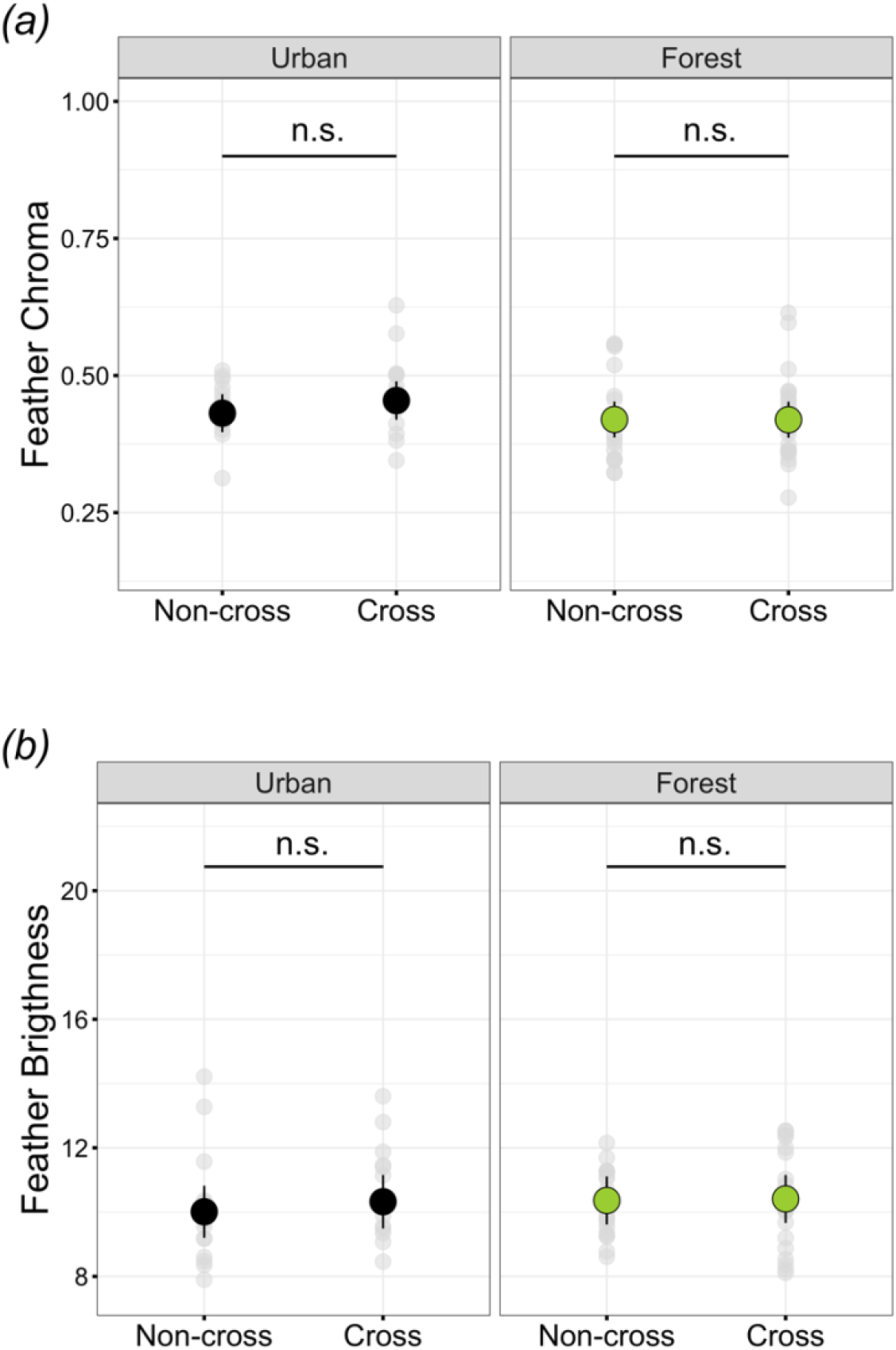
Variation in breast plumage colouration in 15 d old nestlings, reared in urban (left) or forest (right) habitat following a partial cross-fostering between habitats (n= 10 nest pairs; 73 nestlings). *(a)* Chroma reflects the amount of pigment in the feathers (i.e., carotenoids), and *(b)* brightness, reflects the structural quality of feathers. Non-cross: same origin and rearing habitat, (17 urban, 20 forest nestlings); Cross: different origin and rearing habitat (15 urban, 21 forest nestlings). Model mean values (solid-coloured circles) with 95% confidence intervals are plotted, along with raw data (grey circles). n.s. = non-significant differences.

### 3.3 Age-dependent habitat differences in colouration

While we found no effects of urbanisation on yellow colouration in nestling great tits, in Malmö, when we investigated plumage colouration after fledging, in juvenile birds and into adulthood, we found that habitat explained a significant proportion of the variation in yellow chroma, but not in brightness, in an age-dependent manner (Figure 3; “*Habitat x Age*” in Table S6). Consistent with the cross-fostering experiment in *3.2*, we found no differences in chroma between urban and forest nestlings (“*Nestling*” in Figure 3a). However, habitat differences emerged with increasing age: first year urban birds (1 yr) had lower yellow chroma than their forest counterparts (post-hoc Tukey test: t130.2= 3.40, *p=0.001*; Figure 3a), though habitat differences disappeared when comparing urban and forest birds of 2+ yr (post-hoc Tukey test: t111.2= 1.10, p=0.272; Figure 3a).

**Figure 3.**
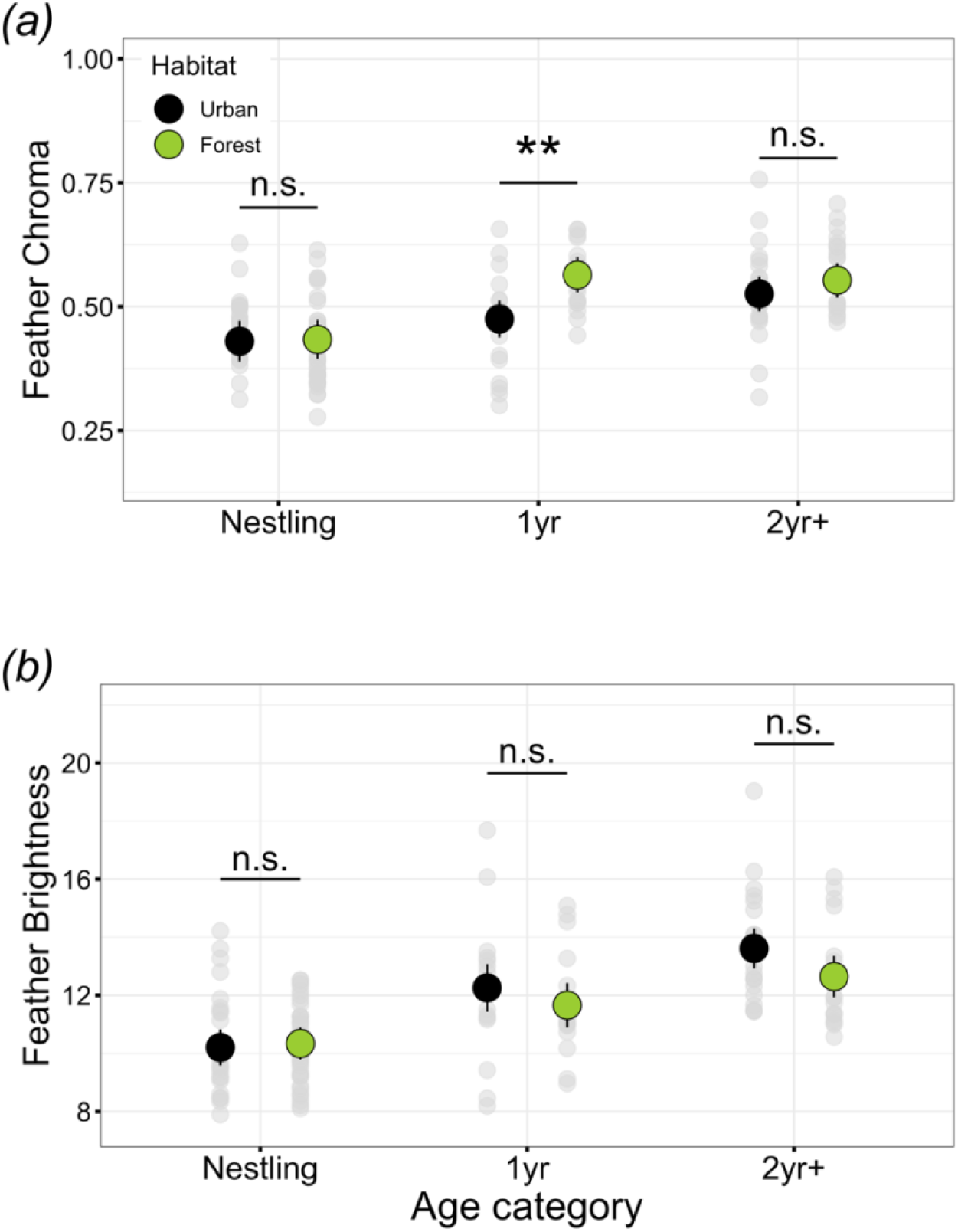
Variation in breast plumage colouration in nestlings and adults in relation to habitat (urban or forest) and age (nestling; first year: 1 yr; and second or more year: 2+ yr) in Malmö, southern Sweden. *(a)* Chroma reflects the amount of pigment in the feathers (i.e., carotenoids), and *(b)* brightness reflects the structural quality of feathers. Note the design is cross-sectional, although nestlings and 1 yr were born in the same year (2013). Sample sizes (urban/forest): nestling (32/41), 1 yr (24/22), 2+ yr (32/41). Urban: black circles; forest: green circles. Model mean values (solid-coloured circles) with 95% confidence intervals are plotted, along with raw data (grey circles). Asterisks denote significant differences between urban and forest habitats with **p< 0.01 (Tukey’s HSD; see table S6 for main effect). n.s. = non-significant differences.

### 3.5 Meta-analysis on colouration differences in response to urbanisation and other anthropogenic disturbances

We performed a meta-analysis on 128 effect sizes from 23 studies that assessed anthropogenic effects (i.e., urbanisation and pollution) on great tit yellow breast colouration traits (Table S2). This revealed that, overall, anthropogenic factors were associated with a decrease in the intensity of plumage colouration of this species (meta-analytic mean: [95%CI] = −0.220 [−0.373, −0.067]; Figure 4a). This model provided evidence for high heterogeneity (*I^2^*_total_ = 94.7%) (Table S7). Geographic location explained 34.0% of the total variation in effect sizes, possibly reflecting among-population variation in the response to anthropogenic disturbances, while 45.5% of the variation was explained by study ID and 15.2% of the total variation was residual variation (Table S7). Further investigation into each of the retrieved plumage traits from the literature supported our findings across the five European regions and, among the evaluated traits, the anthropogenic effect on great tit breast coloration is strongest for chroma (Chroma: −0.213 [−0.330, −0.096]; Figure 4b; Table S7). Interestingly, our meta-analysis also reveals that the observed negative anthropogenic effects on colouration were similar in nestlings and adults (Figure 4c; Table S7). Negative anthropogenic effects on colouration were of a similar magnitude for both urbanisation and pollution, although there is higher uncertainty for pollution, and the effect was not significant (Urbanisation: −0.195 [−0.371, −0.019], Pollution: −0.158 [−0.382, 0.066]; Figure 4d; Table S7). Adjusting our initial meta-analysis estimate to account for potential small-study effects still provided support for an overall negative anthropogenic effect on great tit coloration (adjusted meta-analytic mean: [95%CI] = −0.267 [−0.431, −0.102]; see Equation 22 in Nakagawa et al. 2022) and revealed a significant positive small-study effect (sampling variance slope: 6.045 [2.568, 9.522]); i.e., the higher the sampling variance, the closer the estimate to zero. We did not find evidence for time-lag effects in the dataset used in our meta-analysis (slope: 0.007 [−0.008, 0.023]).

**Figure 4.**
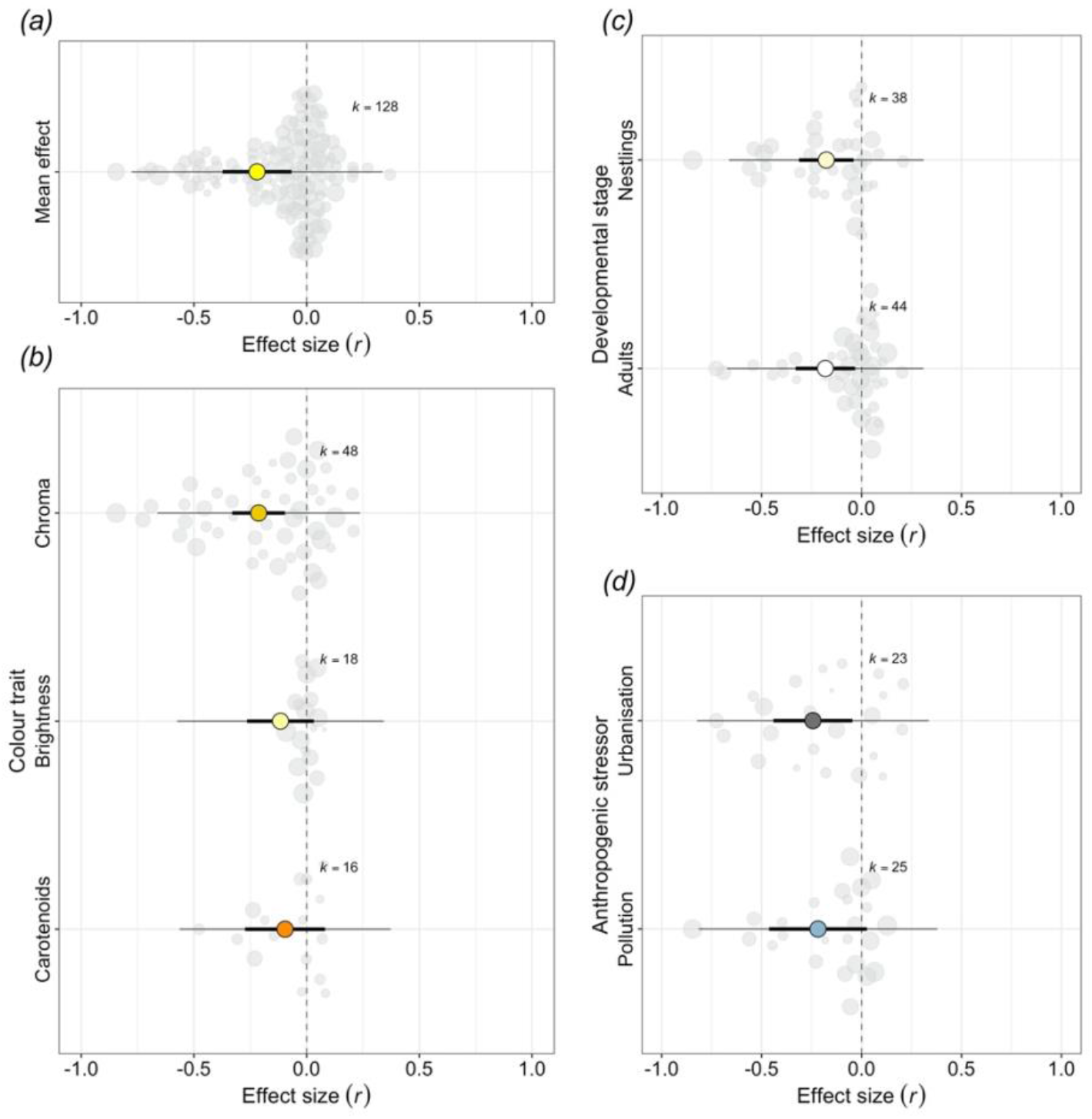
Effect sizes from meta-analysis of the anthropogenic effects on great tit breast plumage coloration. (*a)* Overall mean effect of urbanisation and pollution on great tit breast feather coloration traits (n = 23 studies; k = 128 effects); *(b)* type of colour trait (feather chroma, feather brightness and feather and plasma carotenoid levels); *(c)* developmental stage (adult versus nestling); and *(d)* type of anthropogenic disturbance (pollution versus urbanisation). In all cases, negative values represent “*duller*” yellow colouration/brightness or lower carotenoid concentration in response to urbanisation or exposure to pollution. *(b-d)* Data only corresponds to effect sizes for chroma, brightness and carotenoid levels (n = 20 studies; k = 82 effects). See Table S3 for a detailed description of each moderator levels. Plots show model means and 95% confidence intervals (thick whisker), 95% precision intervals (thin whisker) and individual effect sizes (grey circles; size is scaled to illustrate the sample size from which they were estimated, e.g., the larger the point, the bigger the sample size). Vertical dashed line drawn at an x=0.

## 4 Discussion

It has been long observed that human actions, in particular urbanisation and pollution, can alter wildlife colouration (e.g., Steward, 1977). However, the current knowledge is mostly derived from single-population or geographically confined studies, and there has been no effort to quantify and investigate the spatial replicability of observed patterns at the intra-specific level. Here, using a well-known wild model system in ecology, evolution, and environmental sciences, we empirically demonstrate the overall significant influence of anthropogenic stressors on plumage yellow colouration, supporting previous claims of the “*urban dullness*” phenomenon. Nonetheless, despite the overall effect of urbanisation on plumage colouration, we also reveal spatial variation among distinct localities, spanning the continent of Europe, in the magnitude of the difference in individual yellow colouration between urban and forest populations.

Across Europe, the yellow breast plumage of urban great tits was, on average, paler (i.e., lower chroma) than forest birds, although the variation between forest populations was large when compared with the variation observed between urban populations. On the contrary, plumage brightness did not differ between urban and non-urban populations, indicating that urban yellow breast feathers do not appear duller because they are dirtier. This disparity in the sensitivity to anthropogenic impacts in plumage colour traits is also highlighted in our meta-analysis and previously described for two other great tit urban populations (Biard et al., 2017). While we found an overall negative effect of anthropogenic disturbances on plumage colouration, among the analysed traits, chroma was the only one showing a significant negative effect. Yellow chroma not only reflects an individual’s circulating carotenoid levels and diet quality during moult (e.g., Biard et al., 2006; Eeva et al., 2008, 2009; Isaksson, Von Post, et al., 2007; Peters et al., 2011) but also the ability to incorporate carotenoids into the feathers (Peneaux et al., 2021).

Changes in colour traits have often been suggested as indicators of pollution exposure in wildlife (Lifshitz & St Clair, 2016). It is well established that certain common urban pollutants, such as cadmium or lead, can reduce carotenoid synthesis in plants (Cenkci et al., 2010; Rai et al., 2005), subsequently reducing bioavailability in invertebrate prey (Eeva et al., 2010; Isaksson & Andersson, 2007); ultimately, this manifests in a paler plumage for birds, such as great tits, that feed predominantly on invertebrates during spring and summer (Eeva et al., 2008). However, lower carotenoid bioavailability does not necessarily reflect the direct action of pollution but the result of changes in phenology and invertebrate biomass in polluted habitats (Eeva et al., 2010; Sillanpää et al., 2008). At the European level, it has been shown that the dynamics of insect prey biomass, in particular caterpillars, differ in timing and spatial distribution between urban and non-urban populations (Jensen et al., 2022; Pollock et al., 2017; Seress et al., 2018). Therefore, a diet low in carotenoids, either due to diet quality or quantity, during plumage development (i.e., either during postnatal development in young or post-fledging/breeding moult in juveniles/adults) could contribute to the observed *dullness* pattern across urban great tits, especially for chicks.

In addition, physiological constraints imposed by a trade-off between investment in carotenoid display *versus* homeostasis (e.g., oxidative balance or immune response, in polluted environments: Hutton & McGraw, 2016; Marasco & Costantini, 2016; Peneaux et al., 2021), could also lead to differences in the display of carotenoid-based colouration (but see Koch et al., 2018). In our study, we did not measure any of these proximate mechanisms, but studies in the same populations suggest higher levels of oxidative stress in great tits in relation to urban environmental pollution (Isaksson et al., 2005; Salmón et al., 2018). Nonetheless, in our meta-analysis, the effect of pollution on yellow breast colouration is not significant (but it is in the expected direction), while the response to urbanisation is significant. Urbanisation is a broad disturbance, and therefore, it is likely that the additive or synergistic effects of urban pollution together with the urban environmental mosaic, which influences carotenoid bioavailability, are both key mechanisms driving the observed differences in plumage colouration.

Our meta-analysis indicates that the *dullness* phenomenon in response to anthropogenic disturbance is similar in nestling and adult great tits, which could suggest that the same mechanism drives the paler phenotype despite the different timing of feather development and moult, respectively, in these two contrasting life stages. However, this pattern contrasts with our cross-fostering study in one focal population pair (Malmö). In this location, we found differences in chroma between urban and non-urban adult populations, but not at the nestling stage. Siblings raised in the city and forest did not differ in either chroma or brightness of the yellow breast plumage, regardless of habitat of origin, which suggests that carotenoid availability was not a constraint during the nestling period, at least not during the studied year; thus, habitat-specific differences in carotenoid metabolism or physiological constraints - as previously suggested (Hõrak et al., 2000) - might not explain the observed habitat differences in colouration (specifically chroma) in 1 yr birds. Rather, it is likely that the habitat differences arise after fledging, i.e., during the post-fledging moult in autumn, and reflect temporal variation in environmental conditions (diet and carotenoid availability) during that period. Although, we cannot discount the possibility that observed differences in chroma between urban and forest 1 yr birds are the result of differences in population demographics (Hõrak et al., 2001). Indeed, in the same population, we have previously found evidence for selective disappearance of “*low quality*” birds between fledging and recruitment (Salmón et al., 2017). Nonetheless, juvenile birds usually moult faster than adults and are less efficient in absorbing or utilising dietary carotenoids (Ferns & Hinsley, 2008; Hill, 2002); while this could lead to paler plumage in birds in their first year, compared with older birds, in both urban and forest habitats, the effect might be exacerbated in the urban environment where carotenoid availability is limited compared to the forest and resources often remain lower after the breeding season (Jensen et al., 2022; Pollock et al., 2017; Seress et al., 2018). Although we found habitat differences in colouration in 1 yr birds, these did not persist among birds of 2 yr or older. Adult birds could outcompete juveniles (e.g., Dingemanse & de Goede, 2004), and secure sufficient resources even if scarce; thus, relaxing the habitat differences in colouration in older birds. Nonetheless, the lack of differences in this age group could easily be blurred by age-related changes, since it is not possible to determine exact age of birds beyond their first year.

In conclusion, our multi-analytical study - combining empirical data and a meta-analysis - exemplify that carotenoid-based colouration is consistently paler in response to urbanisation in a model system in avian urban ecology, and paler colouration is likely driven by reduced bioavailability of carotenoids in urban habitats. Yet, spatial and temporal differences are expected across and within populations as the result of the urban landscape heterogeneity (Szulkin et al., 2020), as well as the variable contribution of certain urban stressors such as pollution. Carotenoid signals are proposed as reliable biomarkers of population health, e.g., in birds (Peneaux et al., 2021), although their link with individual quality is often species-specific (Weaver et al., 2018). Despite some attempts to establish the nature of the relationship between carotenoids (and other pigment colouration) and individual quality (Hõrak et al., 2001; Rodewald et al., 2011), we still lack replicated studies linking plumage and individual performance in urban scenarios; such studies are essential to understand the eco-evolutionary implications of the “*urban dullness*” phenomenon.

## Supporting information

Suppementary material

## Acknowledgements

We thank everyone who assisted during the field work and provided logistical support, especially, Sweden: Andreas Nord; Madrid: the ringers from La Herrería for providing the samples from outside Madrid city and Jose Ignacio Aguirre de Miguel; Milan: Michelangelo Morganti and the Lago di Pusiano ringers; Lisbon: Vítor Encarnação and Pardal family. Also, we thank Claire Doutrelant for access to the spectrophotometer.

## Authors’ contributions

D.L-I., P.S. and H.W. conceived the study. C.I., J. P-T., P.S. and H.W. collected the feather samples. D.L-I. analysed the feather colouration. The data analysis was carried out by P.S., P.C-L. conducted the meta-analysis, assisted by D.L-I. and P.S. P.S. and H.W. drafted the initial manuscript with input from all other authors.

