## Supplementary material for "Urbanisation impacts plumage colouration in a songbird across Europe: evidence from a correlational, experimental, and meta-analytical approach": Suppementary material

Full-text articles excluded, with reasons
(n = 8)

Full-text articles assessed for eligibility
(n = 30)

Records excluded
(n = 57)

Records screened
(n = 87)

Records after duplicates removed
(n = 87)

Additional records identified through other sources
(n = 6)

Records identified through WoS searching
(n = 81)

### **Identification**

### **Screening & eligibility**

Studies included in quantitative synthesis (meta-analysis)
(n = 22 + the current study)

### **Included**

**Figure S1.** Preferred Reporting Items for Systematic Reviews and Meta-Analyses (PRISMA) flow diagram for identification and inclusion of studies in the systematic review and meta-analysis. We present the number of papers identified through keyword database searching in addition to records identified through other sources (identified by the authors). Our meta-analysis included effect sizes from the 22 studies identified through the literature search together with the effects sizes arising from the new empirical data presented in this manuscript.

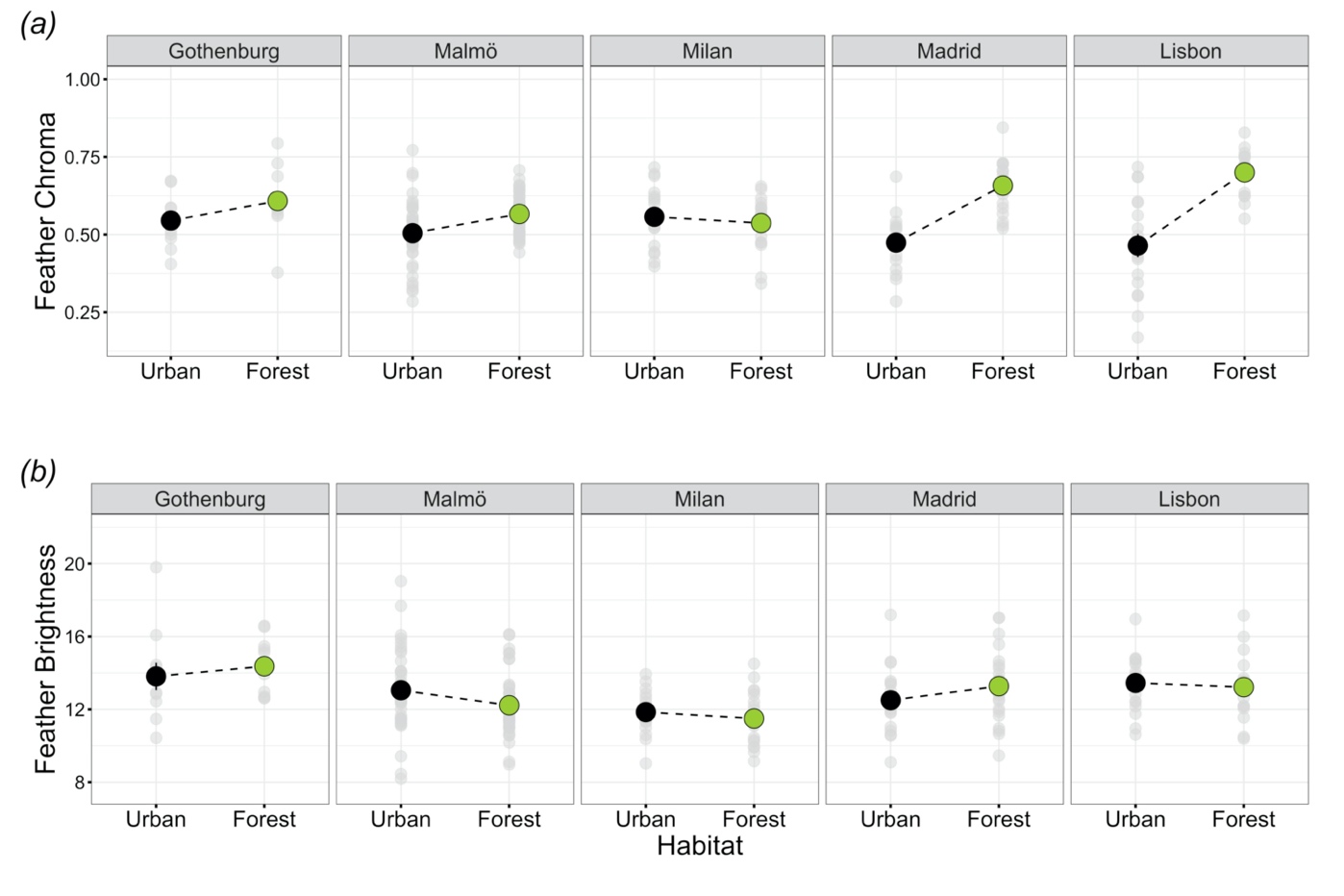
**Figure S2.** Variation in breast plumage colouration (mean ± se of the raw data) among adult great tits (1 yr and 2+ yr) in 5 urban and forest population pairs. *(a)* chroma, reflects the amount of pigment in the feathers (i.e., carotenoids), *(b)* brightness, reflects the structural quality of feathers. Sample sizes (urban/forest): Gothenburg (11/11), Malmö (42/41), Milan (16/18), Madrid (16/20), Lisbon (18/16). The grey individual points in the background show the raw data while the solid-coloured circles represent the mean per population. Note that we found an overall habitat effect across the five populations in chroma but not brightness (see Figure 1; results section and Table S4).

Table S1. Summary of sampling locations: Regional location of urban/forest pairs (city name), year of sampling, season of sampling, centred geographical coordinates of each urban and forest location, urbanisation scale (PC_Urb_, -Principal Component analysis of the land use- positive values indicate higher urbanisation intensity; for details see Salmón, *et al* 2021), number of adult individuals measured in each location (*n*), and distance between the urban and rural location pairs. Cities are sorted in alphabetical order. *Birds captured using mist nets.

|  |  | |  | |  | | *Urban* | | |  | | *Forest* | | |  |
| --- | --- | --- | --- | --- | --- | --- | --- | --- | --- | --- | --- | --- | --- | --- | --- |
| **City** | | **Year** | | **Season** | | **Coordinates** | | **PC_Urb_** | ***n*** | | **Coordinates** | | **PC_Urb_** | ***n*** | **Distance (km)** |
| Gothenburg | | 2015 | | Breeding | | 57°41'24.0"N  11°56'24.0"E | | 2.13 | 11 | | 57°30'00.0"N  12°00'36.0"E | | -2.39 | 11 | 25 |
| Lisbon | | 2014 | | *Post-Breeding | | 38°44'24.0"N  9°10'48.0"W | | 0.65 | 18 | | 38°51'36.0"N  8°49'48.0"W | | -1.86 | 16 | 33 |
| Madrid | | 2014 | | *Breeding | | 40°26'24.0"N  3°43'48.0"W | | 1.60 | 16 | | 40°34'12.0"N  4°09'36.0"W | | -2.05 | 20 | 39 |
| Malmö | | 2014 | | Breeding | | 55°36'00.0"N  12°59'24.0"E | | 2.32 | 42 | | 55°39'00.0"N  13°34'12.0"E | | -2.44 | 41 | 37 |
| Milan | | 2014 | | *Post-Breeding | | 45°31'48.0"N  9°12'36.0"E | | 1.49 | 16 | | 45°49'12.0"N  9°17'24.0"E | | -2.24 | 18 | 34 |

**Table S2.** Summary table of the systematic review of studies linking breast plumage colouration and anthropogenic disturbances, including direct or indirect actions, e.g., pollution, urbanisation, or habitat fragmentation, on great tits (*Parus major*) and used in the meta-analysis. The search was carried out on 2^nd^ August 2021 in six different databases using the following string: TS = ('urban*' OR 'pollut*') AND ('color*' OR 'colour*' OR 'carot*') AND ('Parus major' OR 'great tit' OR 'tit').

| **Reference** | **Location(s)** | **Type of study** | **Colour trait(s)** | **Develop. stage** |
| --- | --- | --- | --- | --- |
| (Biard et al., 2017) | Paris and Niort (France) | 2 urban vs forest populations | Chroma and Brightness | Nestlings |
| (Dauwe & Eens, 2008) | Antwerp area (Belgium) | Heavy metal pollution gradient | Chroma and Hue | Adults |
| (Eeva et al., 1998) | Harjavalta smelter (Finland) | Heavy metal pollution gradient | Colour intensity (using a 1-6 scale) | Nestlings |
| (Eeva et al., 2008) | Harjavalta smelter (Finland) | 1 heavy metal polluted vs control area | Chroma | Nestlings |
| (Eeva et al., 2009) | Harjavalta smelter (Finland) | 1 heavy metal polluted vs control area | Chroma | Nestlings |
| (Eeva et al., 2012) | Harjavalta smelter (Finland) | 1 heavy metal polluted vs control area | Carotenoid concentration in plasma | Nestlings and adults |
| (Eeva et al., 2014) | Harjavalta smelter and Ruissalo (Finalnd) | 1 heavy metal polluted vs control area | Chroma | Nestlings |
| (Geens et al., 2009) | Antwerp area (Belgium) | Heavy metal pollution gradient | Chroma, Brightness and Hue | Nestlings and adults |
| (Giraudeau et al., 2015) | Barcelona (Spain) | Heavy metal pollution levels | Chroma, Brightness and Hue (as a PCA) | Adults |
| (Grunst, Grunst, Pinxten, & Eens, 2020) | Antwerp area (Belgium) | Noise pollution gradient | Chroma, Brightness, Hue and UV Chroma | Nestlings |
| (Grunst, Grunst, Pinxten, Bervoets, et al., 2020) | Antwerp area (Belgium) | Heavy metal pollution gradient | Chroma, Brightness, Hue and UV Chroma | Adults |
| (Hõrak et al., 2000) | Tartu (Estonia) | 1 urban vs forest populations | Chroma | Nestlings |
| (Hõrak et al., 2001) | Tartu (Estonia) | 1 urban vs forest populations | Chroma and Hue | Adults |
| (Hõrak et al., 2004) | Tartu (Estonia) | 1 urban vs forest populations | Hue | Adults |
| (Isaksson et al., 2005) | Gothenburg (Sweden) | 1 urban gradient (urban, semiurban and forest) | Chroma | Nestlings and adults |
| (Isaksson et al., 2006) | Gothenburg (Sweden) | 1 urban vs forest populations | Chroma and UV | Nestlings and adults |
| (Isaksson, McLaughlin, et al., 2007) | Gothenburg (Sweden) | 1 urban vs forest populations | Chroma, Hue and carotenoid concentration in plasma | Nestlings and adults |
| (Isaksson, Von Post, et al., 2007) | Gothenburg (Sweden) | 1 urban vs forest populations | Carotenoid concentration in feathers | Adults |
| (Koivula et al., 2011) | Harjavalta smelter (Finland) | 1 heavy metal polluted vs control area | Feather "yellowness" | Nestlings |
| (S. Ruiz et al., 2016) | Harjavalta smelter and Ruissalo (Finalnd) | 1 heavy metal polluted vs control area | Carotenoid concentration in plasma | Nestlings |
| (S. R. Ruiz et al., 2017) | Harjavalta smelter (Finland) | 1 heavy metal polluted vs control area | Carotenoid concentration in plasma | Nestlings |
| (Sillanpää et al., 2008) | Harjavalta smelter (Finland) | 1 heavy metal polluted vs control area | Carotenoid concentration in plasma and feathers | Nestlings |
| Present study | Malmö, Gothenburg, Milan, Madrid and Lisbon (Sweden, Italy, Spain and Portugal) | 5 urban vs forest populations | Chroma and Brightness | Nestlings and adults |

**Table S3.** Description of moderators used in the meta-analytic models testing for the association between anthropogenic disturbance (i.e., urbanisation and pollution) and great tit breast plumage colouration and carotenoids. k= number of effect sizes per moderator category or level.

| Moderator | Category (k) | Description |
| --- | --- | --- |
| Colouration trait | Chroma (48)  Brightness (18)  Carotenoids (16) | Chromatic component of the yellow colouration, i.e., colouration dependent on the amount of carotenoids deposited in the feather.  Achromatic component of the colouration, i.e., colouration dependent on the feather structure.  Carotenoid concentration in plasma, i.e., concentration of lutein and zeaxanthin, which are directly associated to the chromatic component of colour. |
| Developmental stage | Nestlings (38)  Adults (44) | Individuals used in the study are nestlings, i.e., individuals during their nest phase and dependent on parental care.  Animals used in the study are fully developed individuals, i.e., over a year old. |
| Anthropogenic disturbance | Urbanisation (31)  Pollution (51) | Effect sizes extracted from studies comparing at least one urban and non-urban population. We do not consider, in this category, those studies based on a single population/metapopulation within e.g., a pollution gradient.  Effect sizes extracted for studies comparing the relationship between colouration and environmental pollution, either directly, e.g., via measurement of pollutant levels in biological tissues/faeces, or indirectly, i.e., via comparing populations/subpopulations exposed to different levels of anthropogenic pollution. |

**Table S4.** Summary of linear mixed models exploring the variation in breast plumage colouration among adult great tits (1 yr or 2+ yr) in relation to habitat (urban vs forest). In all models, the sampling location (n= 10 levels) and region (n=5 urban/forest pairs) are included as random effects. Dependent variables were Z-transformed to have mean = 0 and SD = 1. Explanatory power is represented as marginal (R^2^_m_) and conditional (R^2^_c_) values.

|  | Estimate | SE | df | F | p-value |
| --- | --- | --- | --- | --- | --- |
| (a) Dependent variable: *Feather chroma* |  |  |  |  |  |
| (Intercept) | 0.40 | 0.25 |  |  |  |
| Habitat (Urban) | -1.16 | 0.36 | 1, 8.06 | 10.69 | **0.011** |
| Sex (Female) | -0.07 | 0.17 | 1, 195.95 | 0.10 | 0.754 |
| Age (2+ yr) | 0.28 | 0.17 | 1, 197.83 | 11.56 | 0.001 |
| Habitat (Urban) x Sex (Female) | 0.06 | 0.24 | 1, 195.95 | 0.07 | 0.790 |
| Habitat (Urban) x Age (2+ yr) | 0.28 | 0.25 | 1, 197.83 | 1.27 | 0.261 |
| R^2^_m_/ R^2^_c_= 0.23/ 0.40 |  |  |  |  |  |
| (b) Dependent variable: *Feather brightness* |  |  |  |  |  |
| (Intercept) | 0.17 | 0.26 |  |  |  |
| Habitat (Urban) | -0.02 | 0.29 | 1, 4.58 | 0.09 | 0.777 |
| Sex (Female) | -0.25 | 0.19 | 1, 196.70 | 4.70 | **0.031** |
| Age (2+ yr) | -0.07 | 0.19 | 1, 198.52 | 0.10 | 0.754 |
| Habitat (Urban) x Sex (Female) | -0.10 | 0.27 | 1, 197.54 | 0.12 | 0.725 |
| Habitat (Urban) x Age (2+ yr) | 0.23 | 0.28 | 1, 198.44 | 0.66 | 0.416 |
| R^2^_m_/ R^2^_c_= 0.03/ 0.20 |  |  |  |  |  |

**Table S5.** Summary of linear mixed models exploring the variation in breast plumage colouration among nestling great tits (15 days old) in a between-habitat cross-fostering experiment in one of the 5 urban/forest population pairs (Malmö). The cross-fostering design allows to separate between habitat of rearing (urban vs forest) and habitat of origin (urban vs forest). In all models, the nest of rearing and the nest of origin are included as random effects. Dependent variables were Z-transformed to have mean = 0 and SD = 1. Explanatory power is represented as marginal (R^2^_m_) and conditional (R^2^_c_) values.

|  | Estimate | SE | df | F | p-value |
| --- | --- | --- | --- | --- | --- |
| (a) Dependent variable: *Feather chroma* |  |  |  |  |  |
| (Intercept) | -0.30 | 0.30 |  |  |  |
| Habitat of rearing (Urban) | 0.51 | 0.41 | 1, 16.75 | 0.84 | 0.371 |
| Habitat of origin (Urban) | -0.01 | 0.19 | 1, 9.33 | 1.04 | 0.333 |
| Sex (Female) | 0.04 | 0.13 | 1, 47.46 | 0.10 | 0.757 |
| Habitat of rearing (Urban) x Habitat of origin (Urban) | -0.30 | 0.24 | 1, 43.57 | 1.64 | 0.207 |
| R^2^_m_/ R^2^_c_= 0.04/ 0.77 |  |  |  |  |  |
| (b) Dependent variable: *Feather brightness* |  |  |  |  |  |
| (Intercept) | 0.01 | 0.28 |  |  |  |
| Habitat of rearing (Urban) | -0.01 | 0.38 | 1, 18.29 | 0.17 | 0.682 |
| Habitat of origin (Urban) | 0.03 | 0.28 | 1, 13.08 | 0.18 | 0.674 |
| Sex (Female) | 0.06 | 0.22 | 1, 57.81 | 0.07 | 0.788 |
| Habitat of rearing (Urban) x Habitat of origin (Urban) | -0.25 | 0.42 | 1, 46.51 | 0.36 | 0.553 |
| R^2^_m_/ R^2^_c_= 0.01/ 0.28 |  |  |  |  |  |

**Table S6.** Summary of linear mixed models exploring the variation in breast plumage colouration across great tit age categories (nestling, first year: 1 yr and second or more year: 2+ yr) in relation to habitat (urban vs forest) in one of the population pairs (Malmö). In all models, the territory location is included as a random effect. Dependent variables were Z-transformed to have mean = 0 and SD = 1. Explanatory power is represented as marginal (R^2^_m_) and conditional (R^2^_c_) values.

|  | Estimate | SE | df | F | p-value |
| --- | --- | --- | --- | --- | --- |
| (a) Dependent variable: *Feather chroma* |  |  |  |  |  |
| (Intercept) | 0.80 | 0.20 |  |  |  |
| Habitat (Urban) | -0.96 | 0.29 | 1, 61.67 | 5.64 | **0.021** |
| Sex (Female) | 0.01 | 0.14 | 1, 90.28 | 0.22 | 0.639 |
| Age (2+ yr) | -0.11 | 0.25 | 2, 119.42 | 19.14 | **<0.001** |
| Habitat (Urban) x Sex (Female) | -1.35 | 0.26 | 1, 90.28 | 0.15 | 0.699 |
| Habitat (Urban) x Age (2+ yr) | 0.08 | 0.20 | 2, 119.42 | 3.22 | **0.044** |
| R^2^_m_/ R^2^_c_= 0.32/ 0.72 |  |  |  |  |  |
| (b) Dependent variable: *Feather brightness* |  |  |  |  |  |
| (Intercept) | 0.14 | 0.21 |  |  |  |
| Habitat (Urban) | 0.25 | 0.31 | 1, 147.00 | 2.85 | 0.094 |
| Sex (Female) | -0.15 | 0.18 | 1, 147.00 | 0.77 | 0.381 |
| Age (2+ yr) | 0.47 | 0.25 | 2, 147.00 | 41.86 | **<0.001** |
| Habitat (Urban) x Sex (Female) | -0.63 | 0.22 | 1, 147.00 | 0.10 | 0.756 |
| Habitat (Urban) x Age (2+ yr) | 0.08 | 0.26 | 2, 147.00 | 1.61 | 0.204 |
| R^2^_m_/ R^2^_c_= 0.37/ 0.37 |  |  |  |  |  |

**Table S7.** Summary of the multilevel meta-analytic mixed-effect models. Bold estimates indicate confidence intervals (CI) that do not overlap zero. *k* is the number of effect sizes per level, and *n* is the number of studies. For the “*intercept-only model*”, i.e., meta-analytic mean, the associated heterogeneity (*I^2^*) explained for each random factor and the whole model is shown. Estimated heterogeneity explained by the moderators (R^2^_marginal:_ R^2^_m_) is also shown.

|  | **Estimate [95% CI]** | ***k*** | ***n*** |
| --- | --- | --- | --- |
| *Intercept-only model* |  |  |  |
| Intercept | **-0.220 [-0.373, -0.067]** | 128 | 23* |
| *I^2^ _location_* = 33.99%  *I^2^ _Study ID_* = 45.47%  *I^2^ _Residual_* = 15.22%  *I^2^ _Total_* = 94.69% |  |  |  |
| *Colouration trait* |  |  |  |
| Chroma | **-0.213 [-0.330, -0.096]** | 48 | 15* |
| Brightness | -0.116 [-0.264, 0.032] | 18 | 5* |
| Carotenoids | -0.096 [-0.274, 0.082] | 16 | 6 |
| R^2^_m_= 4.89% |  |  |  |
| *Developmental stage* |  |  |  |
| Nestlings | **-0.177 [-0.313, -0.040]** | 38 | 16* |
| Adults | **-0.181 [-0.331, -0.032]** | 44 | 10* |
| R^2^_m_= 0.009% |  |  |  |
| *Type of anthropogenic disturbance* |  |  |  |
| Urbanisation | **-0.195 [-0.382, -0.018]** | 31 | 8* |
| Pollution | -0.158 [-0.3824, 0.066] | 51 | 12 |
| R^2^_m_= 0.05% |  |  |  |

*Includes the present study.
